# Experimental tests of the quantum property of protein folding

**DOI:** 10.1101/044230

**Authors:** Liaofu Luo

## Abstract

Experimental tests on the quantum property of protein folding are discussed. It includes: the test of the instantaneousness of torsion transition through observation of protein structural change in a short time scale of microsecond; the test of non-Arrhenius temperature dependence of protein folding rate and other biomolecular conformational changes; and the search for the narrow spectral lines of the protein photo-folding.

---

Protein folding kinetics is usually studied by use of molecular dynamics (MD) simulation which is based on the classical mechanics [1]. However, when we observe the folding event at the molecular level, the classical approach is simply too limit to realize a complete solution of the folding mechanism problem. It is well known that fluorescence and phosphorescence are phenomena closely related to protein folding and such phenomena can only be understood in the context of quantum mechanics. Thus, the application of quantum theory to protein folding should be more reasonable and logically simple. On the other hand, despite the remarkable achievements made in 1960 in developing quantum biochemistry of nucleic acid molecules, the quantum theory past only studied the motion of electrons in a macromolecule but cannot discuss the conformational change that is related to the variables of molecular shape. Recently, we proposed a quantum theory on protein folding [2][3], or more generally, on macromolecular conformational change. As is well known, the molecular shape is fully determined by a set of coordinates of bond length, bond angle and torsion angle. Among these variables the torsion is slow and others are fast. The torsion energy is the lowest in all forms of biological energies, even lower than the average thermal energy. The torsion angles are easily changed even at physiological temperature. Following Haken’s synergetics, the slow variables always slave the fast ones. By the adiabatic elimination of fast variables we obtained the Hamiltonian describing the conformational transition of protein. Finally we deduced a formula for protein folding rate in analytical form. It is worth mentioning that this formula can automatically include solvent effects since this effect has been considered by the modified torsion potential. A long-standing problem is why the protein folding rate always exhibits the curious non-Arrhenius temperature dependence [4]. Our formula can explain all these experiments well [5]. Furthermore, we have succeeded in applying this theory to study RNA folding and deduced a general statistical relation between folding rate and chain length for both protein and RNA molecules [6].

However, it is argued that quantum theory is applicable to macromolecules. A popular viewpoint is: due to the environmental quantum entanglement the decoherence makes quantum picture of molecule ineffective, and the estimation of decoherence time seems indicate that the molecule with large mass moves in a classic trajectory [7]. But this estimate is very rough. We found that if the same model is applied to the torsion degrees of freedom for a protein, the decoherence time will be long enough [2]. Therefore one may also conclude that the concept of quantum folding is still correct. So, the final solution of the problem on what rules, quantum or classical, are obeyed by the macromolecular conformational motion needs to be awaited only after more direct experimental evidences.

This is an important experiment. If the test has a positive result, then firstly, the boundary between classical and quantum physics will be modified and the applicability of quantum mechanics will expand to the slow variable degrees of freedom of the macromolecule; secondly, many strange phenomena of sudden change that is of great importance in molecular biology and molecular genetics will can be explained from the idea of quantum transition.

In summary, what should be cut away by Occam’s Razor in studying conformational change of biological macromolecule ‐‐ classical or quantum? Following our point the quantum transition provides a unifying and logically simple theoretical starting point. We have developed a quantum theory on the shape transition of macromolecule. Now we will suggest several experiments to test the quantum property of the conformational motion.

## 1 Test of the instantaneousness of the torsion transition through observation of protein structural change in a short time scale of microsecond

Whether the protein folding is quantum or classical can be directly tested from the instantaneous nature of quantum transition. It is widely accepted that the change of electronic state in a atom is quantum transition and the transition is instantaneous. The instantaneousness is characteristic of the quantum transition. So the puzzling problem on whether the molecular torsion transition is quantum or classical can also be solved from the observation of the instantaneous change of the torsion angle.

Because the timescale of protein folding is microsecond to millisecond we suggest to observe the detailed change of torsion angle Δ_*i*_*φ*_*i*_, Δ_*i*_*ψ*_*i*_, Δ_*i*_*χ*_*i*_ versus folding time *t* in a time duration of folding, where φ and ψ represent the backbone dihedral angle, χ the side chain dihedral angle and the subscript *i* means the *i*-th residue. The curve of Δ_*i*_*φ*_*i*_, Δ_*i*_*ψ*_*i*_, Δ_*i*_*χ*_*i*_ versus *t* is continuous in classical theory. However for quantum folding, when the concentration of protein molecules is low enough that the single-molecule behavior could be observed, we will see a special picture of fluctuation, namely, the continuous curve changes to several points randomly scattered with time in the duration of protein folding. Note that in the quantum description when one says the folding rate 1/ τ. it does not mean the folding continued τ seconds but refers only to transition 1/ τ times per second on average.

**Fig 1.**
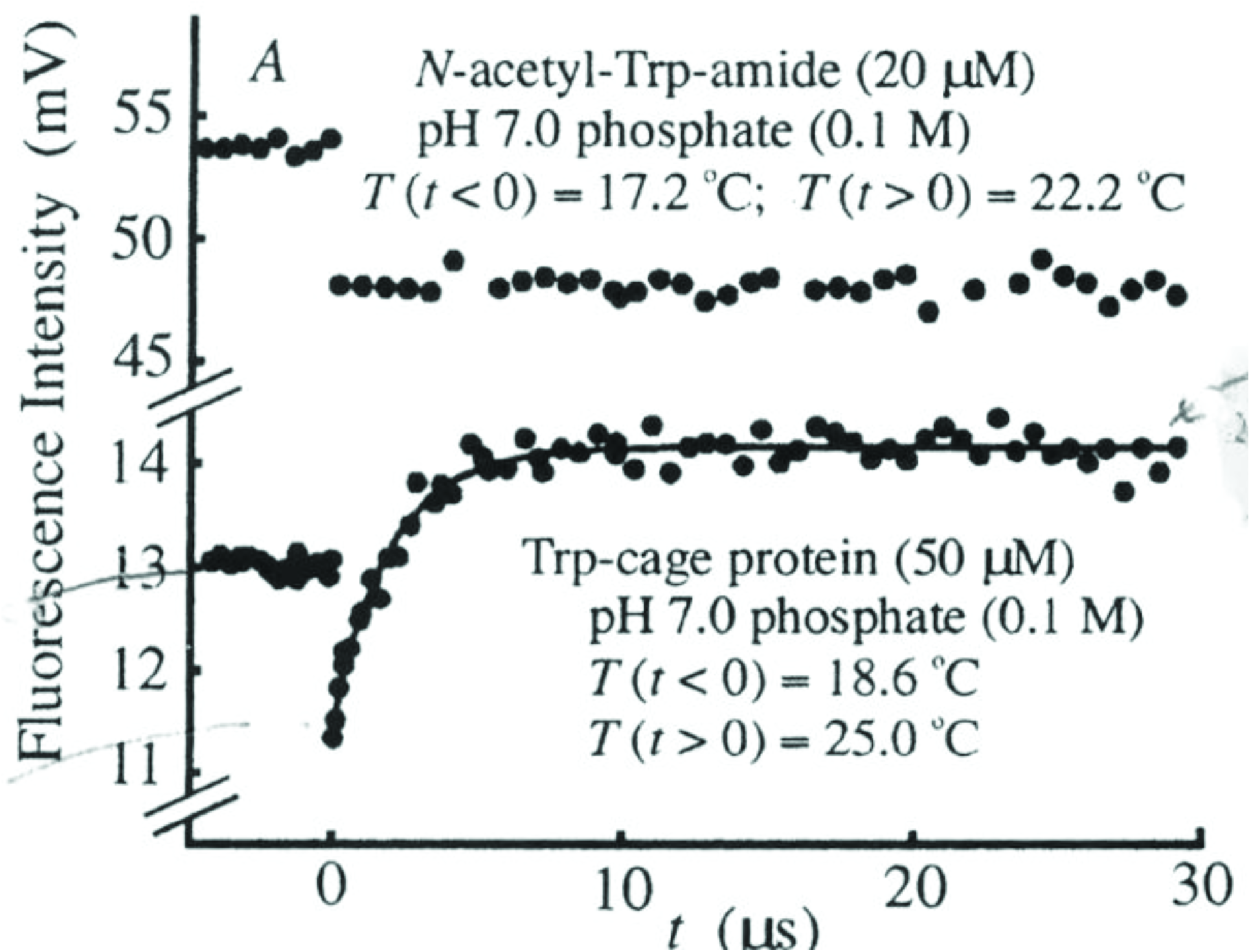
20-Residue Trp-Cage Folds. Fluorescence response to 6.4 °C T-jump of Trp-cage and response of free Trp(upper trace) as a reference. (after Qiu et al, JACS 2002 [8])

Qiu et al used laser temperature-jump spectroscopy to measure the folding rate of the 20-residue Trp-cage protein, found the fluorenscence intensity (FI) increasing rapidly from 11.5mV to 14 mv in 4 μ s, and determined the rate 4 μ s.[8] Is the folding/unfolding event occurring in the 4 μ s in accordance with the classical physical law or with the law of quantum physics? Consider the case of the gradual decrease of molecular concentration. In the beginning, the fluorenscence intensity will be weakened accompanying with the lowering Trp-cage concentration, but the shape of the FI-t curve remains not changed. As the concentration decreased to very low, the fluctuation appears. For quantum folding, the torsion angle takes only two possible values corresponding to folding and unfolding states respectively. The Trp fluorenscence can only be measured in unfolding state. So, in the duration of 4 μ s the fluorenscence randomly appears and each occurrence corresponds to one unfolding event. However, if the folding / unfolding obeys the laws of classical physics the torsional angle changes continuously and the Trp fluorescence can be recorded only when the protein reaches the unfolding state. Thus the fluorescence will be measured near the end of 4 microsecond unfolding process. Two pictures are different from each other. We suggest to make this observation of fluorenscence fluctuation to test the folding obeying classical or quantum.

Femtosecond stimulated Raman spectroscopy (FSRS) is a new ultrafast spectroscopic technique that provides vibrational structural information with high temporal (50-fs) and spectral (10-cm^‒1^) resolution [9]. The time resolution 50-fs is enough to study the detailed structural changes in protein folding. Taking some time points during protein folding to make FSRS measurement, recording the vibration spectrum information and observing its time evolution one can follow the details of the conformational change. This may provide another method to study the problem on whether the protein folding is classical or quantum.

## 2 Study of the temperature dependence of protein folding and other biomolecular conformational changes

The non-Arrhenius temperature dependence of folding rate cannot be explained by MD [1]. However, from quantum folding theory we have deduced the temperature dependence of protein folding rate (*W*) as follows

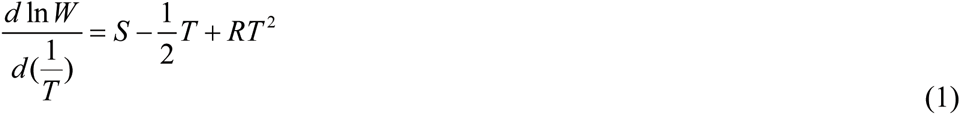

The non-linearity of Eq (1) means the non-Arrhenius temperature dependence. The particular temperature dependence reflects the quantum properties of the folding mechanism. We have proved the theoretical Eq (1) is in accordance with the experimental temperature dependence of the folding rate for all proteins whose experimental data were available [5]. This provides a strong evidence on quantum property of protein folding. The temperature dependence (1) comes mainly from the torsional transition (the slow-variable degree of freedom) of the molecule and basically irrespective of the fast-variable. We have considered two models of fast variable for protein folding: the fast variables are stretching-bending of bonds and the fast variables are electron motion. Both models deduce the basically same temperature dependence, namely Eq (1) for stretching-bending and replacement of 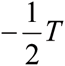 by 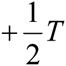 in (1) for electrons as fast variables [2].

Moreover, we have studied the quantum histone modification in gene expression [10], the quantum glucose transport across membrane [11], the common step of RNA folding from compact intermediates [6] and the pluripotency conversion of genes in stem cell [12]. All these processes are “slaved” by torsional transition and they should exhibit the basically same temperature dependence. Tests of temperature dependence give check points for the quantum property of the conformational change.

To explore the fundamental physics behind the folding more deeply and clarify the quantum nature of the folding mechanism more clearly we have studied the protein photo-folding processes, namely, the stimulated photon emission or absorption in protein folding and the inelastic scattering of photon on protein (photon-protein resonance Raman scattering).[13] We have proved that the stimulated photo-folding rates and the photon-protein resonance Raman scattering sections show the same temperature dependence as protein folding. The experimental tests of these points are awaited for.

## 3 Search for extra-narrow spectral lines of protein photo-folding

Due to the coupling between protein structure and electron motion, the electronic transition will inevitably lead to the structural relaxation or conformational changes of the protein. Therefore the spectrum of protein photo-folding includes information of several kinds of quantum transitions: the electron energy-level transition, the transition between vibrational energy-levels of the molecule, the transition between rotational energy-levels of the molecule and the transition between different molecular conformations. Conformational transitions are somewhat like the rotational transitions, but the rotational transition refers to the whole molecule, while the conformational transition is related only to the dihedral angle rotation of local atomic groups.

Because of the coupling with vibration and torsional transition the electronic spectrum is broadened and the spectral band is formed. The width of the spectral band is determined by the torsion vibration frequency, in the order of 10^13^ sec^-1^, one hundredth or thousandth of the electronic transition frequency. The spectral band includes a large amount of spectral lines from the transitions between torsion-vibration states in different conformations. The transition is a kind of forbidden transition since the overlap integral between initial and final torsional wave functions is very small, about 10^-5^. Therefore, in protein photo-folding we shall observe each spectral line of electronic transition broadened to a band which includes abundant extra-narrow spectral lines. The width of the latter is five orders smaller than the natural linewidth.[13] This is an important prediction of quantum folding theory.

From the experimental point of view, to observe the extra-narrow spectral line a high-precision and high-resolution spectroscopy is needed. The spectral resolution of FSRS is 10-cm^‒1^, corresponding to Δ*v* = 3×10^11^*s*^‒1^ [9]. This resolution is already close to the range of the width that the spectral line of ultra narrow conformational transitions can be searched.

## Acknowledgment

The author is indebted to Dr Jun Lv for his numerous helpful discussions.

